# Bio-Docklets: Virtualization Containers for Single-Step Execution of NGS Pipelines

**DOI:** 10.1101/116962

**Authors:** Baekdoo Kim, Thahmina Ali, Carlos Lijeron, Enis Afgan, Konstantinos Krampis

**Affiliations:** Center for Translational and Basic Research and Belfer Research Building, Hunter College of The City University of New York, New York; Department of Biological Sciences, Hunter College of The City University of New York, New York; Johns Hopkins University, Department of Biology, Baltimore, MD; Department of Physiology and Biophysics, Institute for Computational Biomedicine, Weil Cornell Medical College, NY

**Keywords:** Docker, Bioinformatics, NGS, RNAseq, ChIPseq

## Abstract

**Background:** Processing of Next-Generation Sequencing (NGS) data requires significant technical skills, involving installation, configuration, and execution of bioinformatics data pipelines, in addition to specialized post-analysis visualization and data mining software. In order to address some of these challenges, developers have leveraged virtualization containers, towards seamless deployment of preconfigured bioinformatics software and pipelines on any computational platform.

**Findings:** We present an approach for abstracting the complex data operations of multi-step, bioinformatics pipelines for NGS data analysis. As examples, we have deployed two pipelines for RNAseq and CHIPseq, pre-configured within Docker virtualization containers we call Bio-Docklets. Each Bio-Docklet exposes a single data input and output endpoint and from a user perspective, running the pipelines is as simple as running a single bioinformatics tool. This is achieved through a “meta-script” that automatically starts the Bio-Docklets, and controls the pipeline execution through the BioBlend software library and the Galaxy Application Programming Interface (API). The pipelne output is post-processed using the Visual Omics Explorer (VOE) framework, providing interactive data visualizations that users can access through a web browser.

**Conclusions:** The goal of our approach is to enable easy access to NGS data analysis pipelines for non-bioinformatics experts, on any computing environment whether a laboratory workstation, university computer cluster, or a cloud service provider,. Besides end-users, the Bio-Docklets also enables developers to programmatically deploy and run a large number of pipeline instances for concurrent analysis of multiple datasets.

## FINDINGS

### BACKGROUND

Analysis of NGS data involves multiple technical steps such as installation of the software components of bioinformatics pipelines; coordinating format conversions and data flow between pipeline components; managing software versions and updates; automating execution for multiple runs; supplying the required computational and data storage infrastructure; and last but not least, providing an intuitive user interface for non-bioinformatics experts. To overcome these challenges, bioinformatics software developers have leveraged technologies such as virtual machines and Docker containers ([1], [2]) for distributing preconfigured bioinformatics software that can run on any computational platform. The use of virtualization saves significant development time and cost, as the software does not need to be set up from scratch at each laboratory. The increased interest for applications of virtualization for NGS data analysis is evident through many recent studies, ranging from comparing performance of virtual machines to conventional computing [3], and bioinformatics-specific Docker container repositories [4].

The Galaxy server [5] provides an innovative approach for deployment of command-line software through an online Graphical User Interface (GUI), and has had a great impact on making NGS data analysis tools and pipelines easily accessible to non-bioinformatics experts. In addition, the Galaxy ecosystem provides the Toolshed [6] for downloading and installing a range of commonly used bioinformatics software, with a workflow composition canvas on the GUI and a high-performance pipeline execution engine in the backend. Standardized workflow descriptions in eXtensible Markup Language (XML) files allows transferring NGS analysis pipelines across Galaxy installations at different laboratories, but the bioinformatics software used in a pipeline needs to be re-installed at each location through the ToolShed or manually.

A number of different virtual machines with the Galaxy server [7] are currently available, but a search for the term “pipeline” through the list returns only two entries. These are easily accessible through VirtualBox [8], but unless users setup VirtualBox shared folders and connect the Galaxy data libraries to these, non-experts will have to upload data through the Galaxy interface and result in duplication within the virtual machine. Furthermore, the Galaxy Docker containers on the list presume a level of software expertise, since users need to manually run them on a local server or on the cloud. A simpler version of the “bread-and-butter” NGS data analysis pipeline implemented in the present study version for RNAseq has been previously published as Galaxy workflows [9], in addition to CHIPseq [10]. Given the computing time limits or storage quota of 250GB on the public Galaxy server [11], the size of current NGS datasets and the amount of intermediate output generated by the bioinformatics tools in these pipelines, it will only be feasible for researchers to perform two or three complete runs of either pipeline under a single account. Alternatively, users can start their own Galaxy compute cluster on the Amazon cloud service through CloudMan [12] but a significant amount of setup steps are required [13]. Nonetheless, even researchers with credit cards linked to research funds might be reluctant to repeatedly pay for leasing computing time and for the costs associated with maintaining data on the cloud, compared to a one-time investment for a laboratory compute server. Finally, a significant bottleneck for non-bioinformatics experts is that the pipeline data output requires additional post-processing, filtering and visualization in order to generate scientific insights and figures that can be included in manuscripts.

### METHODS

In this study, we present the Bio-Docklets virtualization containers by combining Docker, Galaxy, and a “meta-script” (Fig. 1a) that enables users to run complex, multi-step data analysis pipelines, as simply as running a single command. In addition, we have implemented Python code (Fig. 1b) that leverages the BioBlend software library [14] to access Galaxy API, and automate pipeline execution through the Galaxy pipeline engine running inside the container. Additional scripts inside the containers (Fig. 1c,d,e) automate retrieval of supporting data such as reference genome sequences and indices, while also initializing the required environment variables within the containers, starting and monitoring the pipeline execution, and saving all generated outputs to the host file system before the container terminates. In the final stage (Fig 1f) we have integrated the pipelines with the Visual Omics Explorer framework (VOE, [15]), through custom code that post-processes the raw pipeline output and loads it on interactive data visualizations. Furthermore, the Bio-Docklets containers expose the server ports where the Galaxy is accessible, so that the full interface of the Galaxy canvas is available should the users need to edit the pipelines.

**Fig. 1.**
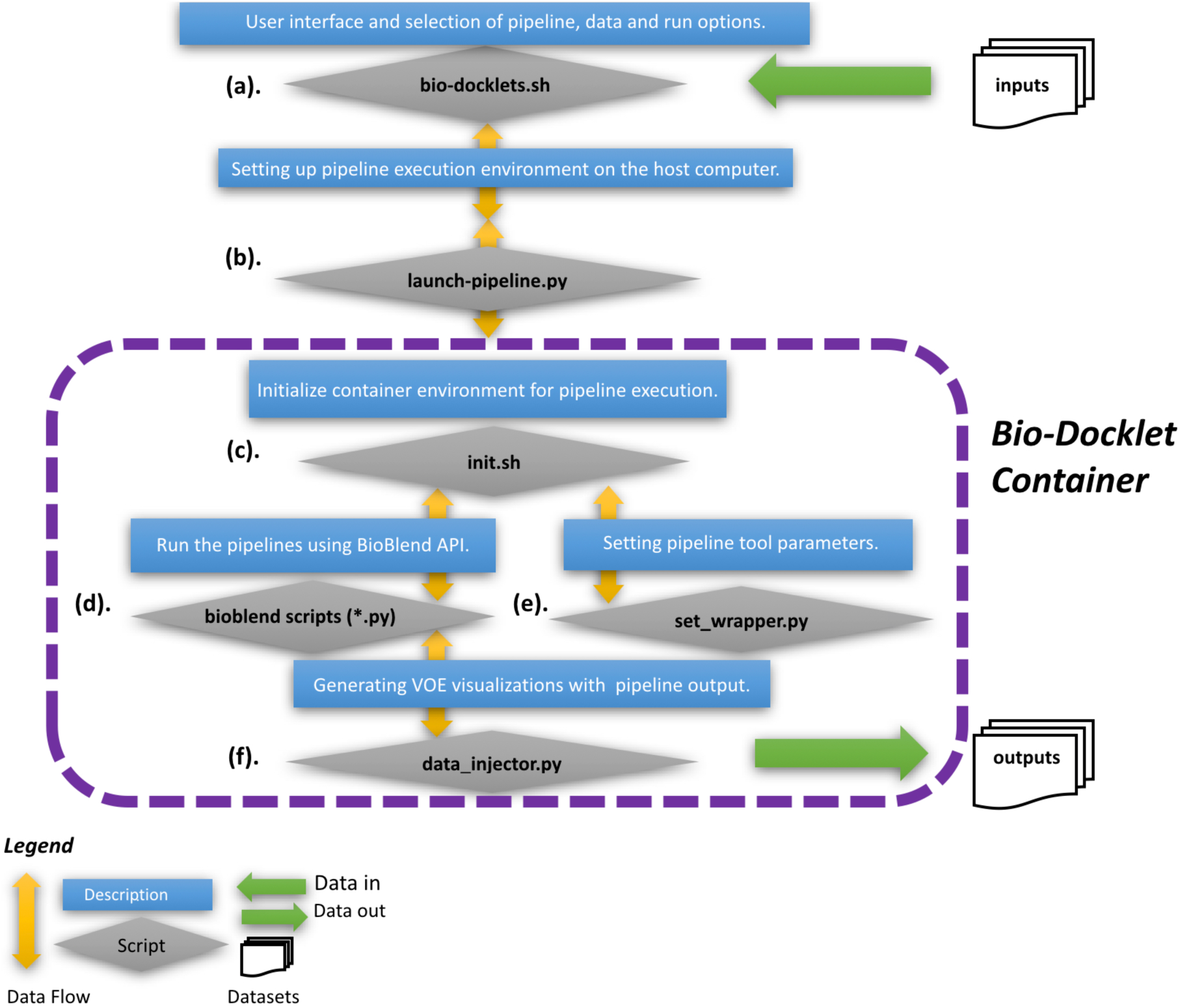
The Bio-Docklets environment with an **(a).** Interactive meta-script to allow users to start the pipelines and set data directories and other parameters for the host system; **(b, c, d, e).** Code that connects to the Galaxy API, automates retrieval of supporting data such as reference genomes, initializes the required environment variables and starts the pipelines, and; **(f).** post-processing of the pipeline output for generating interactive visualizations with the results, that can be accessed independently of the containers and exported for use in manuscripts.

For our implementation, we started from a base Docker container, where we first installed Galaxy and then created two identical copies, the first for implementing the RNAseq [16] pipeline and the second for CHIPseq [17]. In each container, we installed the bioinformatics tools used in the pipelines from the Galaxy Toolshed, or manually if not available. The Galaxy workflow canvas was then used for composing the pipelines (Fig. 2a, 2b), and upon completion and testing, the containers were published on a DockerHub repository [18]. Next, we implemented the “meta-script” that interactively guides the users to select which pipeline to run (Fig 2c), prepares the input and output file directories, verifies that the user's data are in the correct format, retrieves supporting data such as referece genome indices, while it also automatically downloads and runs the Bio-Docklets containers from the repository, in addition to installing the Docker virtualization layer if not present on the host computing system (**Suppl.**). All data generated from the pipelines are saved to the output directory specified by the user, in addition to the VOE visualization files that are in HTML / Javascript-D3 [19] format. These can be simply opened in a web browser independently of the Bio-Docklets, as ready-to-use, interactive visualizations and also be exported as high-resolutions, manuscript-ready graphics.

**Fig. 2.**
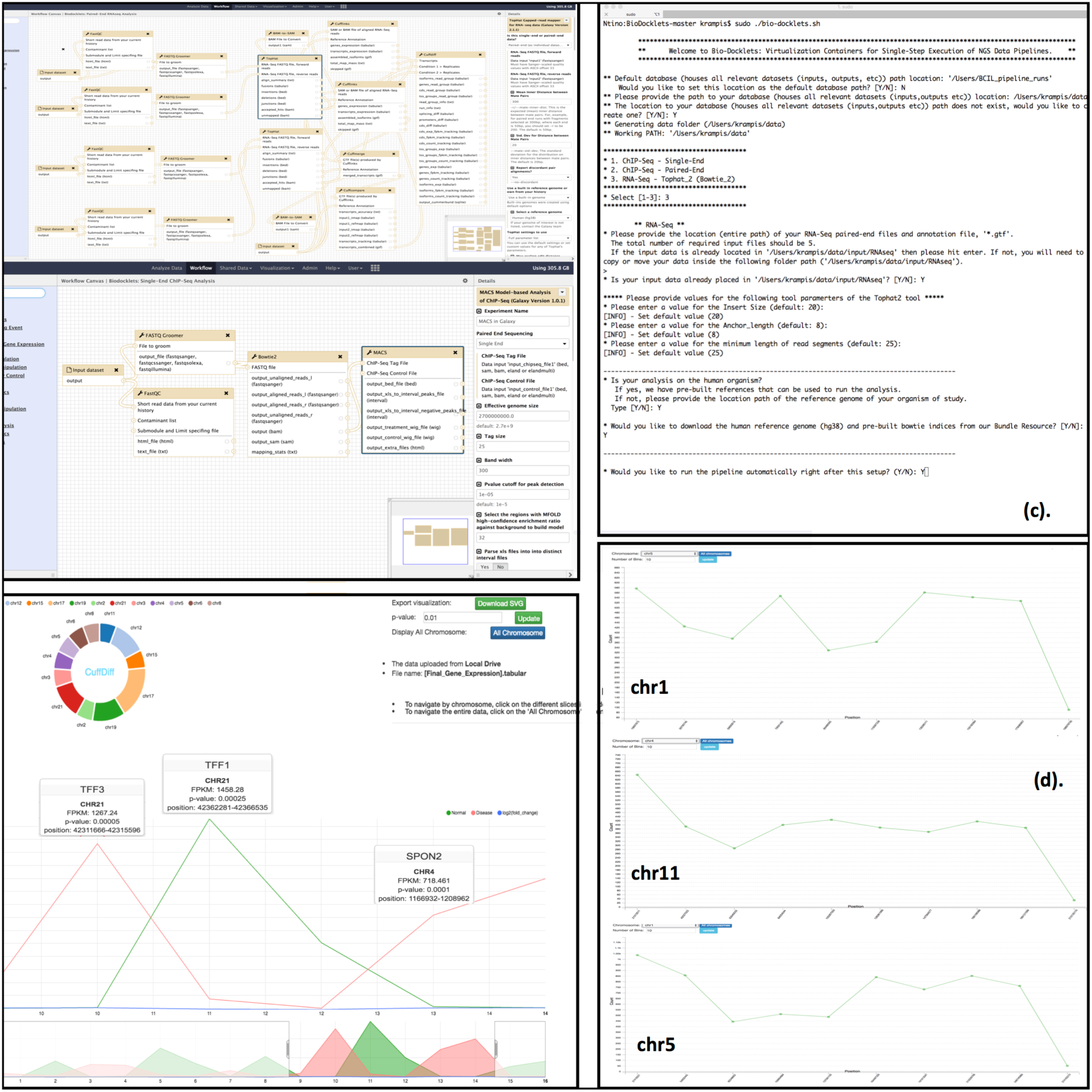
**(a,b).** Galaxy workflow canvas running inside the Bio-Docklets, with the composed RNAseq and CHIPseq pipelines respectively; **(b).** User interface of the “meta-scipt”, that interactively guides the users to select which pipeline to run, input and output file directories and reference genome indices; **(c).** Post-processed pipeline output, loaded on interactive HTML / Javascript-D3 visualizations using the Visual Omics Explorer framework that can be opened in a web browser and also exported as high-resolutions, manuscript-ready graphics.

Our target audience is research teams that do not have any bioinformatics expertise, but are generating NGS data using sequencing technology such as Illumina MiSeq or MiniSeq [20]. The Bio-Docklets approach aims to help these groups perform a basic analysis and interpretation of their datasets with minimal effort. Regarding computational infrastructure, existing laboratory computers with at least 4 CPU cores and 500GB disk storage capacity, can provide enough computational capacity to run the containers and process the approximately 30 million reads generated per run by these instruments [21], or researchers scale up to 200 million reads with a larger capacity server (Table 1) as presented in the next section.

**Tabel 1:**
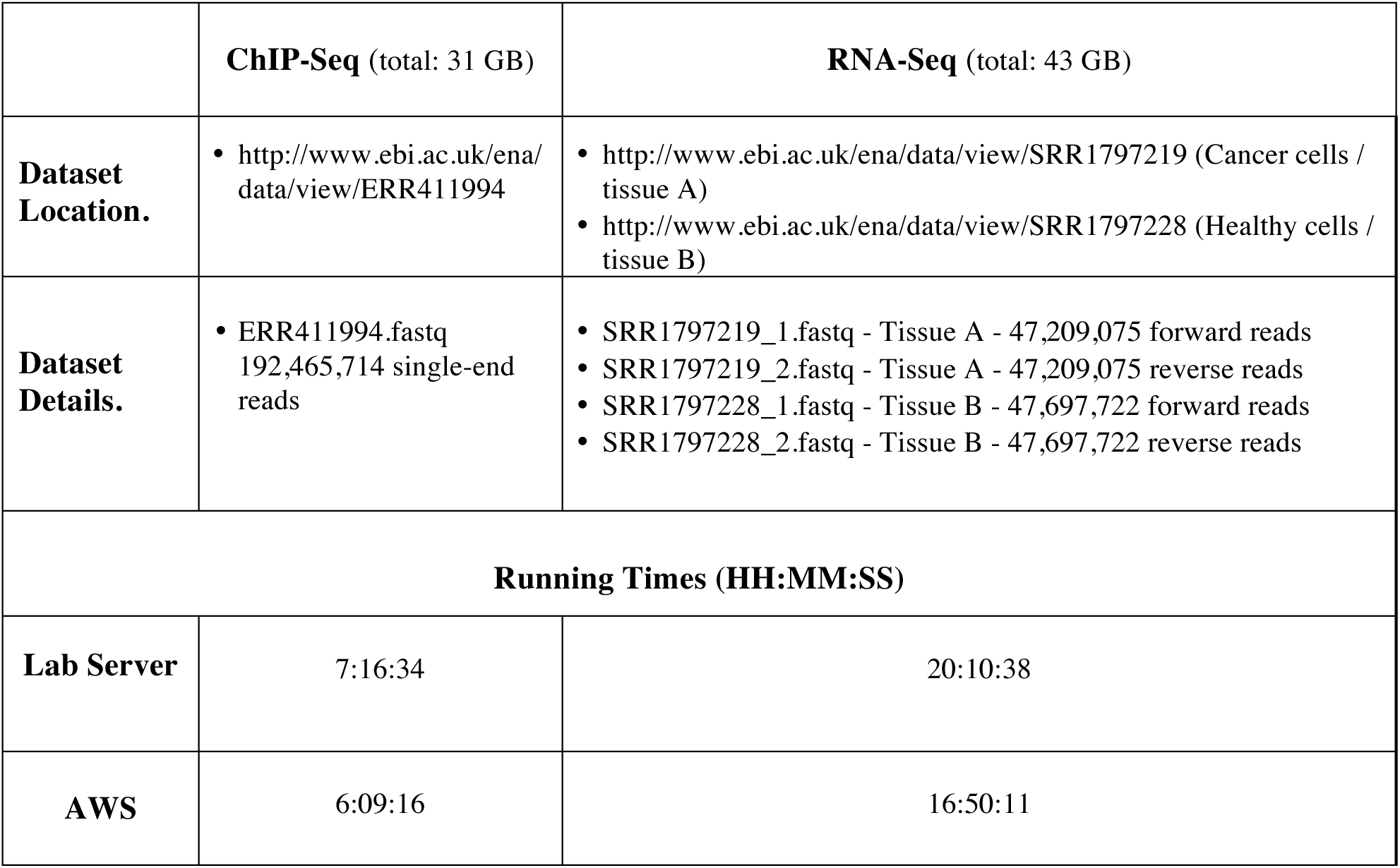
Benchmark run times of the Bio-Docklet pipeline containers with the ChIPseq and RNAseq pipelines, using as input large-scale NGS data downloaded from public databases.

### PERFORMANCE AND TESTING

In order to test the computational performance and functionality of the Bio-Docklet containers, we used publicly available NGS data from the European Bioinformatics Institute archive (EBI, http://www.ebi.ac.uk/). First, we processed a dataset with approximately 190 million Acute Myeloid Leukemia (AML) single-end reads through the CHIPseq Bio-Docklet, provided as input to the container a file of 31GB size (EBI reference ERR411994, Table 1). The RNAseq Bio-Docklet was tested with 43GB of input data (EBI reference SRR1797219 and SRR1797228), containing a total of 188 million reads (47 million x 4, given two paired read files, for both the cancer and healthy tissue samples, Table 1). We run each Bio-Docklet in turn on our lab computer server (32GB RAM, 4 CPU Intel Xeon), measuring a total running time of 20 hours and 10 minutes for RNAseq to complete, while for CHIPseq the time was significantly lower at 7 hours and 16 minutes (Table 1). This was expected, given the lesser computational requirements for the alignment of single-end reads in the CHIPseq dataset. We also processed the same datasets by running the Bio-Docklets on a compute server with more computational capacity (Table 1), which was rented from the Amazon Web Services cloud (http://aws.amazon.com). In the CHIPseq output we observed a large number of significant peaks (p-value < .001) on chromosomes 1, 4, 5, 7, 8, 11, 16 and 19 (Fig 2), which harbor histone interactions with active role to tumor genesis [22], while for RNAseq we found differentially expressed genes that are active regulators in cancer progression [23].

### DISCUSSION

Currently, a number of other bioinformatics software development projects are utilizing Docker virtualization, including BioShaDock that provides a curated repository of pre-built bioinformatics containers, BioContainers / BioDocker [24] that implements an aggregator and search engine across Docker repositories, and BioBoxes [25] defining a standardized interface for running bioinformatics tools pre-installed in containers. Using the search terms “Galaxy” and “pipeline” returned 4 and 34 entries for BioShaDock, 8 and 30 for BioContainers respectively, while BioBoxes currently includes a total number of 8 containers. The former two, provide a great way for developers in the bioinformatics community to distribute tools and pipelines and reach the right audience, given that from our experience DockerHub is a very large repository and specialized bioinformatics containers might be missed in the keyword search results. Given than BioShaDock and BioContainers provide containers that are “Automatic Builds” from Dockerfiles, no citation or other information how to run the pipelines is provided on the site, and we had to perform an online search for the website of the original bioinformatics project, in order to find the documentation. On the other hand, BioBoxes provide a standardized interface where users can run a bioinformatics tools with a single command that includes input and output data directories, but there is no interactive script or other menus to guide the users through the different parameters. BioBoxes introduces a novel approach for standardizing the pipeline implementation that needs to be adopted by bioinformatics developers, and to the best of our understanding does not include multi-step pipeline execution capabilities or a workflow engine.

The NGSeasy [26] project implements a modular approach where the bioinformatics components used in each step of the pipeline are on different Docker containers, and the pipeline run is coordinated by a single “master” container that runs all components based on a workflow specification in a file provided by the user. While significant software development in NGSeasy abstracts the pipeline run and coordination among the different containers, users are still required to manually install Docker and setup the required data directories, while there is no straightforward method for providing runtime parameters for the algorithms used in the pipeline. Two example projects leveraging Docker in combination with a graphical user interface, are GUIDock [27] and BioDepot-workflow-Builder (BwB, [28]). The former focuses on providing pre-configured containers with specialized tools such as CytoScape [29], but in order to access the graphical interface users are required to install Xquartz [30] and other specialized components, which can be challenging for non-technical users. The BwB suite provides a pipeline composition canvas, similar to an open-source alternative of the Seven Bridges platform [31] in addition to separate Docker containers for each pipeline component. However, significant software development expertise is needed for uers to write their own widgets and install the tools. On the other hand, our approach using Galaxy allow you to easily download and install tools available from the ToolShed, without any software development effort. Starting a Galaxy instance on the cloud requires significant technical expertise [32].

In this study, we have abstracted complex bioinformatics data operations in a format that is fully portable across computational platforms, by encapsulating pre-configured NGS pipelines within virtualization containers we call Bio-Docklets. Our goal is to enable researchers to perform advanced genomic data analysis without any prior technical expertise, by running multi-step data pipelines as simply as running as a single bioinformatics tool. Through the use of virtualization and the Galaxy workflow engine, the Bio-docklets approach provides bioinformatics “black-boxes” that expose a single input and output endpoint, while internally perform complex data workflow operations. Furthermore, the BioBlend API in combination with the code included in the Bio-Docklets enables developers to programmatically manage data inputs, output, and control the Galaxy workflow engine that runs the pipelines, in order to build bioinformatics infrastructures with a large number of container instances for parallel analysis of multiple datasets. As an alternative, we have also considered lightweight workflow engines such as NextFlow (http://www.nextflow.io), but settled on Galaxy given that the ToolShed provided easy installation for some of the components we included in the pipelines. Furthermore, the on-screen information printed by our meta-script provides users with access to the Galaxy server and workflow canvas running in the Bio-Docklets, which users can access on their web browser in order to edit the pipelines. For a future update, we are working towards implementing a software platform where users can author Bio-Docklets by composing pipelines through the Galaxy running in the containers, and then automatically publish on the bioinformatics container repositories or DockerHub for broad access by the community.

## Declaration

### Ethics (and consent to participate)

Not applicable

### Consent to publish

Not applicable

### Authors' contributions

BK developed all automated script and constructed method architecture. TA implemented method pipelines, performed data analysis and method validation. CL assisted with method validation and manuscript preparation. KK wrote the manuscript, supervised all the work from conception to manuscript preparation and review.

***Availability of data and materials***

### List of abbreviations used (if any)

**NGS** Next Generation Sequencing **GUI** Graphical User Interface **XML** eXtensible Markup Language **RNAseq** RNA sequencing **ChIPseq** Chromatin Immunoprecipitation sequencing **AWS** Amazon Web Services **HTML** HyperText Markup Language **VOE** Visual Omics Explorer **USB** Universal Serial Bus **API** Application Programming Interface

## ACKNOWLEDGEMENTS

### Funding

Supported by the CTBR and RCMI grant from NIMHD (G12 MD007599) and WCMC-CTSC (2UL1TR000457-06). The authors would like to thank all members of the Bioinformatics Core Infrastructures and Krampis’ Lab for their feedback during manuscript preparation.

